# *wnt4a* promotes female development and reproductive duct elongation in zebrafish

**DOI:** 10.1101/421362

**Authors:** Michelle E. Kossack, Samantha K. High, Rachel E. Hopton, Yi-lin Yan, John H. Postlethwait, Bruce W. Draper

## Abstract

In laboratory strains of zebrafish, sex determination occurs in the absence of a typical sex chromosome and it is not known what regulates the proportion of animals that develop as male or female. Many sex determination and differentiation genes that act downstream of a sex chromosome are well conserved among vertebrates, but studies that test their contribution to this process have mostly been limited to mammalian models. In mammals, WNT4 is a signaling ligand that is essential for ovary and Müllerian duct development, where it function, in part, to antagonize the male-promoting FGF9 signal. Wnt4 is highly conserved in non-mammalian vertebrates, but it is not known if Wnt4 plays a role in sex determination and/or the differentiation of sex organs outside of mammals. This is an especially interesting question in teleost, such as zebrafish, because they lack an Fgf9 ortholog. Here we show that *wnt4a* is the ortholog of mammalian *Wnt4,* and that *wnt4b* was present in the last common ancestor of humans and zebrafish, but was lost in mammals. We found that *wnt4a* is expressed in the somatic cells of juvenile gonads during the time sex determination likely occurs. We show that *wnt4a* loss-of-function mutants develop predominantly as males and conclude that *wnt4a* activity promotes female sex determination in zebrafish. Additionally, both male and female *wnt4a* mutants are sterile because their reproductive ducts do not connect to the vent, where *wnt4a* is normally expressed. Yet when dissected from homozygous *wnt4a* mutant gonads, both sperm and eggs can produce fertile offspring. Together these results strongly argue that Wnt4a is a conserved regulator of female sex determination and reproductive duct development in non-mammalian vertebrates.

**SUMMARY:** Wnt4 is a key regulator of ovary development in mammals, but it is not known if it plays a similar role in other vertebrates. Here we show that zebrafish *wnt4a* is the ortholog of mammalian *Wnt4*. We show that *wnt4a* is expressed in zebrafish somatic gonad cells during the time sex determination likely occurs. Through analysis of *wnt4a* mutants, we show that Wnt4a promotes female sex determination and the development of the male and female reproductive. We conclude that Wnt4/Wnt4a is likely a conserved regulator of ovarian and reproductive duct development in all vertebrates

## INTRODUCTION

Zebrafish (*Danio rerio*) is a major model research organism, yet little is known about its underlying molecular mechanism of sex determination. Zebrafish that were domesticated for laboratory use do not have a single sex chromosome; instead, several loci have been identified that appear to influence sex ratios in a strain-dependent manner (Bradley *et al*. 2011; Anderson *et al*. 2012; Howe *et al*. 2013). In contrast, non-domesticated strains use a ZZ/ZW genetic sex determination mechanism, with the major sex locus being located on chromosome 4 (Wilson *et al*. 2014). Until this locus is characterized, the conserved genes involved in sex determination and/or differentiation in other vertebrates may offer insight into zebrafish sex determination.

Although domesticated zebrafish do not possess a major sex determining locus, some progress has been made toward understanding how sex is determined. Overt sex differences are not apparent until about 20-30 days post-fertilization (dpf), when males and females tend to differ in number of oocytes, with female gonads generally having more oocytes than male gonads (Wang *et al*. 2007). It is therefore presumed that definitive sex determination occurs sometime between 20-25dpf, though an earlier time point cannot be ruled out.

Prior to sex determination, the zebrafish gonad, like the mammalian gonad, is bipotential. Starting around 10dpf, a subset of germ cells in all zebrafish enter meiosis to form early stage oocytes (Takahashi 1977) establishing the bipotential gonad. At the same time, based on gene expression analysis, the somatic gonad is a mixture of male- and female-like cells. For example, we and others have shown that during this stage, some somatic gonad cells begin to express the female-specific gene *cyp19a1a* (*aromatase*), while neighboring cells express the male-specific gene *anti-Müllerian hormone* (*amh*) (Rodríguez-Marí *et al*. 2005; Leerberg *et al*. 2017). Beginning around 20dpf, oocytes in some individuals undergo apoptosis as these gonads begin the transition to testis development. In contrast, oocytes in gonads destined to become ovaries continue their maturation (Uchida *et al*. 2002; Maack and Segner 2003). Importantly, if all germ cells, or specifically oocytes, are ablated prior to or during the bipotenital phase, all animals develop as phenotypic males (Slanchev *et al*. 2005; Siegfried and Nüsslein-Volhard 2008; Rodríguez-Marí *et al*. 2010; Dai *et al*. 2015). These results led to the model that oocytes produce a signal that stabilizes female development; in the absence of a threshold level of the signal, the animals develop as males. The identity of the oocyte-producing signal or how it affects sex determination of somatic gonad cells remains to be determined.

Growing evidence suggests that Wnt signaling may also play a conserved role in teleost sex determination and/or differentiation. In zebrafish, over-expression of a dominant-negative TCF transcription factor, the downstream effector of the canonical Wnt signaling pathway, increases the production of males over females (Sreenivasan *et al*. 2014). Thus canonical Wnt signaling appears to be involved in female sexual development in zebrafish. In mammals, WNT4 is the WNT ligand involved in sex determination (Vainio *et al*. 1999), but the specific Wnt ligand that functions to regulate sex determination in zebrafish remains to be determined.

WNT4 (wingless-type MMTV integration site family, member 4) is a signaling ligand that binds to the Frizzled receptor and activates the canonical Wnt signaling pathway (as reviewed in (Nusse and Clevers 2017)). In mammals, which utilize an XX/XY genetic sex determination system, Wnt4 is critical for female sex determination. In addition the early gonad is bipotential and both sexes initially express the male-specific gene *fibroblast growth factor 9* (*Fgf9*) in the overlying gonadal epithelium while the underlying mesonephros expresses the female-specific gene *Wnt4* (Vainio *et al.* 1999; Bowles *et al*. 2010). In the absence of *Sry*, the Y-linked male sex determinant, WNT4 inhibits the expression of FGF9 and the gonad develops into an ovary that continues to express WNT4. In contrast, expression of SRY in XY animals stabilizes *Fgf9* expression, which in turn leads to the inhibition of W*nt*4, and thus testis development (Kim *et al*. 2006). Importantly, XX mice lacking WNT4 sex-revert to male (Vainio *et al*. 1999), demonstrating that WNT4 is necessary for female development. Interestingly, simultaneous loss of WNT4 and FGF9 in XY animals results in normal testis development, arguing that the main role of FGF9 in males is to antagonize the female-promoting WNT4 signal (Jameson *et al*. 2012). Additionally, mutations in the human *WNT4* gene can lead to a variety of reproductive diseases that affect ovary development, including Polycystic ovary syndrome (PCOS) (Pellegrino *et al*. 2010) and female <u>se</u>x reversal and dysgenesis of <u>k</u>idneys, <u>a</u>drenals, and lungs (SERKAL), where chromosomally XX gonads lacking WNT4 function no longer develop as an ovary and instead develop as testicular tissue (Mandel *et al*. 2008).

Here we show that Wnt4a functions to promote female sex determination in zebrafish. In addition, we show that Wnt4a is required for the development of the reproductive ducts in both male and female zebrafish. These results therefore demonstrate that Wnt4 is a conserved regulator of female sex determination or differentiation in a non-mammalian vertebrate.

## METHODS

### Phylogenetic analysis

Phylogenetic analysis of Wnt4a was conducted as previously described (Vilella *et al*. 2009) using the compara gene tree tool found at ensemble.org.

### Zebrafish rearing

The IACUCs at the University of California Davis and the University of Oregon approved all animals used in this study (protocols #18483 and #14-08R, respectively). Zebrafish husbandry was performed as previously described (Westerfield 2007) with the following modifications to the larval fish (5-30dpf) feeding schedule: 5-12dpf: 40 fish/250mL in static fish water (4parts/thousand (ppt) ocean salts) were fed rotifers (*Brachionus plicatilis*, L-type) twice daily *ad libitum*. 12-15dpf: 40 fish/one liter gently flowing fish water (<1ppt ocean salts) were fed both rotifers and freshly hatched *Artemia* nauplii *ad libitum* twice daily. 15-30dpf: 40 fish/one liter gently flowing fish water (<1ppt ocean salts) were fed freshly hatched *Artemia* nauplii *ad libitum* twice daily.

### Fish lines

The ziwi:EGFP transgenic line and *wnt4a(fh294)/+* mutant line were developed in an AB background. The *wnt4a(fh294)/+* mutant line was created by treating adult AB zebrafish males with ENU and identifying sequence changes in the *wnt4* gene (Moens 2009). The resulting *wnt4a* mutation is a nucleotide substitution that creates a premature stop codon at amino acid 307 of 352 (Moens 2009). The *wnt4a(uc55)* and *wnt4a(uc56)* alleles were produced by CRISPR/Cas9 genome editing, with the following guide RNA targeting exon two: 5’ – AGCTGTCGTCGGTGGGGAGC(PAM)-3. *wnt4a(uc55)* and *wnt4a(uc56*) are predicted to cause a translational frame shift in exon two.

### Fluorescent in situ hybridization (FISH)

The *wnt4a in situ* probe was generated by PCR (see Supplemental Table 1 for primers) producing a 1987bp fragment. These were cloned into pGEM-T Easy vector (Promega). For whole mount studies on 10-30dpf gonads, *wnt4a* was hybridized at a concentration of 1:200 at 65°C to permeabilized tissue for 48 hours, after which whole-mount gonads were developed using an Alkaline-Phosphatase reaction with FastRed (Sigma) for 8 hours. Vasa antibody (1:1500) staining was performed after a glycine wash as described in (Draper 2012). Gonads were imaged with an Olympus FV1000 laser scanning confocal microscope. Acquired images were adjusted equally using ImageJ.

### RT-PCR for genes of interest

RNA was extracted from the gonads of three individuals at 90dpf or 30dpf and RNA was combined before reverse transcription. Amplification of *wnt4a, wnt4b, cyp19a1a, amh,* and *rpl13a* was performed with the following program: Step 1: 94°C for two minutes, Step 2: 94°C for 15 seconds, 65°C for 15 seconds, 72°C for 15 seconds, repeated 28 times, Step 3: 72°C for two minutes. Primers listed in Supplemental Table 1. Products were run on a one percent agarose gel and imaged.

### Genotyping

#### wnt4a (fh294) PCR

The primers used for genotyping the *wnt4a*(fh294) mutant line are listed in the Supplemental Table 1 using the following PCR conditions: Step 1: 94°C for one minute; 35 cycles of: 94°C for 30 seconds, 60°C for 30 seconds, 72°C for one minute and 30 seconds; followed by 15°C until program was ended. Resulting amplicons were digested with DdeI at 37°C overnight (Moens 2009). Sizes of bands after DdeI digest: 384bp (wild-type), 270bp + 114bp (mutant).

#### wnt4a(fh295) PCR

Genomic DNA was extracted and fh295 mutant fish were identified using HRMA (Dahlem *et al*. 2012) using primers listed in Supplemental Table 1. The program was as follows: Step 1: 95°C for 1 minute; Step 2: 39 cycles of 94°C for 10 seconds, 69°C for 15 seconds; Step 3: 94°C for 20 seconds; Step 4: 72°C for 20 seconds; Followed by a melt profile from 80-92°C with increments of 0.2°C.

#### wnt4a(uc55 and uc56) PCR

The primer pairs used for PCR genotyping are listed in Supplemental Table 1, using the following PCR protocol: Step 1: 94°C for one minute; Step 2: 34 cycles of: 94°C for 10 seconds, 55°C for 10 seconds; Step 3: 72°C for 15 seconds. The PCR products were separated on a three percent agarose gel: wild-type: 123bp, *wnt4a(uc55)*: 140bp, w*nt4a(uc56)*:149bp.

### Sex Ratios and Characterization of Mutant Phenotypes

At 90dpf or more, fish were genotyped and sacrificed. Secondary sexual characteristics were examined, and the gonad of each fish was dissected to confirm gonadal sex. A subset from animals of each genotype was randomly measured for standard length. Characterization of mutant development was performed by Anti-Vasa antibody staining as described previously (Draper 2012).

### Mutant fertility assessment

*wnt4a*(uc55) and *wnt4a*(fh294) heterozygous fish were set up in a crossing cage with either heterozygous or mutant counter parts. Eggs were collected and counted for percent fertilization. Following mating tests, *wnt4a*(uc55) and *wnt4a*(fh294) heterozygous or mutant males and females were squeezed for sperm or eggs following techniques described in (Walker and Streisinger 2007). If eggs or sperm were not released, gonads were dissected and fertilized with the heterozygous counterpart *in vitro*. Percent of eggs successfully fertilized and embryo survivability was tracked to five dpf.

### Hematoxylin and Eosin staining

At 90dpf, *wnt4a(uc55)* and *wnt4a(fh295)* mutant and wild-type fish (n=3) were identified by PCR genotyping, were euthanized, and fixed in Bouin’s fixative for 24hrs. The samples were then embedded in paraffin, cut into 7µm sections and stained with Hematoxylin and Eosin (H&E). Reproductive ducts were examined and representative images were taken at 5X on a Zeiss Axiophot microscope.

### Micro Computed Tomography (microCT)

At 90dpf fish were genotyped and confirmed to be wild-type or *wnt4a(uc55)* mutant (n=3). Fish were anesthetized with MS22 for 5 minutes and exsanguinated by cutting off the tail and placing the fish head up in a filter column in a 1.5mL micro-centrifuge tube followed by centrifuging at 40rcf at room temperature for five minutes. The blood clot was then removed and the fish was centrifuged again. Fish were then fixed in PFA for at least 24 hours. Before imaging, fish were placed in 2.75% Lugol’s, (14mL/fish) for 24 hours and then washed in 1x PBS for one hour.

Zebrafish were imaged at the Center for Molecular and Genomic Imaging (UC Davis) with X-ray CT. 90dpf fish were embedded in one percent agar gel and positioned in a 15mm diameter conical tube. The straw was mounted on an aluminum post for placing in the CT scanner. X-ray tomographic images were obtained on the Center’s MicroXCT-200 specimen CT scanner (Carl Zeiss X-ray Microscopy). Samples were mounted on the scanner’s sample stage, which can be positioned to the submicron level. Scan parameters were adjusted based on the manufacturer’s recommended guidelines. The 4x objective of the MicroXCT was chosen for optimal spatial resolution of reproductive ducts. The source and detector distances were set at 30mm and 10mm, respectively. Once the source and detector settings were established, the optimal x-ray filtration was determined by selecting one of 12 proprietary filters; in this case, no filtration was necessary. Following this procedure, the optimal voltage and power settings were determined for optimal contrast (80kV and 100µAmp). 1600 projections over 360 degrees were obtained with 0.75 seconds per projection. The camera pixels were binned by two and the source-detector configuration resulted in a voxel size of 5.0693µm. Tomographic images were reconstructed with a center shift (7.11 pixels) and beam hardening parameter value of 0.2 to obtain optimized images. A smoothing filter of kernel size 0.7 was applied during reconstruction. Images were reconstructed into 16-bit values.

### In situ Hybridization on sections

Animals were collected at multiple stages of zebrafish male reproductive duct development. Animals were then euthanized, fixed, and cryosectioned as previously described (Rodríguez-Marí *et al*. 2005). The probe for *wnt4a* was created using primers listed in Supplemental Table 1. The *wnt4a* cDNA was cloned using the TOPO vector and used to synthesize DIG-labeled probes. For *in situ* hybridization experiments, two 25dpf, two 35dpf male, and two 55dpf male zebrafish were used.

### Data and reagent availability

All fish lines are available upon request, and will be deposited at the Zebrafish International Stock Center (ZIRC). All supplementary figures are available at Figshare:

FigS1 RT-PCR analysis of *wnt4a* expression in males and females (.tif).

FigS2. *wnt4a* is expressed in somatic gonad cells (.tif).

FigS3. CRISPR/Cas9-induced *wnt4a* mutations (.tif).

FigS4. External phenotype of *wnt4a* mutants (.tif).

FigS5. Histological analysis of *wnt4a* mutant gonads (.tif).

FigS6. Movie, 3D renderings of the male reproductive ducts (.mov)

FigS7. Movie, 3D renderings of the female male reproductive ducts (.mov)

FigS8. Time course of reproductive duct extension in wild-type and *wnt4a* mutants.

## RESULTS

### *wnt4a* is the ortholog of mammalian *Wnt4*

Mammalian genomes contain a single *WNT4* gene but most teleost genomes have two *Wnt4-*related genes called *wnt4a* and *wnt4b* (Ungar *et al*. 1995; Liu *et al*. 2000). Connectivity of teleost genomes to the human genome requires accurate designation of orthologs, which necessitates an understanding of gene histories. The two teleost *wnt4-*related genes could have resulted from either: 1) gene duplication after the divergence of mammalian and teleost lineages, for example, in the teleost genome duplication event (Amores *et al*. 1998; Postlethwait *et al*. 1998; Jaillon *et al*. 2004) or 2) duplication before the divergence of the human and zebrafish lineages followed by loss in the mammalian lineage. To test these models, we studied gene phylogenies and conserved syntenies. Phylogenetic analysis showed that ancestral lobe-finned vertebrates had two *wnt4-*related genes, the *wnt4a* and *wnt4b* clades, because several lobe-finned animals (birds, reptiles, and coelacanth) have these genes today (Figure 1A). Ancestral ray-finned vertebrates also had both *wnt4* clades because orthologs of both appear in spotted gar and teleost fish (Figure 1A). This evidence shows that the last common ancestor of human and zebrafish had both *wnt4a* and *wnt4b,* ruling out the hypothesis that *wnt4a* and *wnt4b* arose in the teleost genome duplication and supporting the loss of *wnt4b* in the origin of mammals.

**Figure 1.**
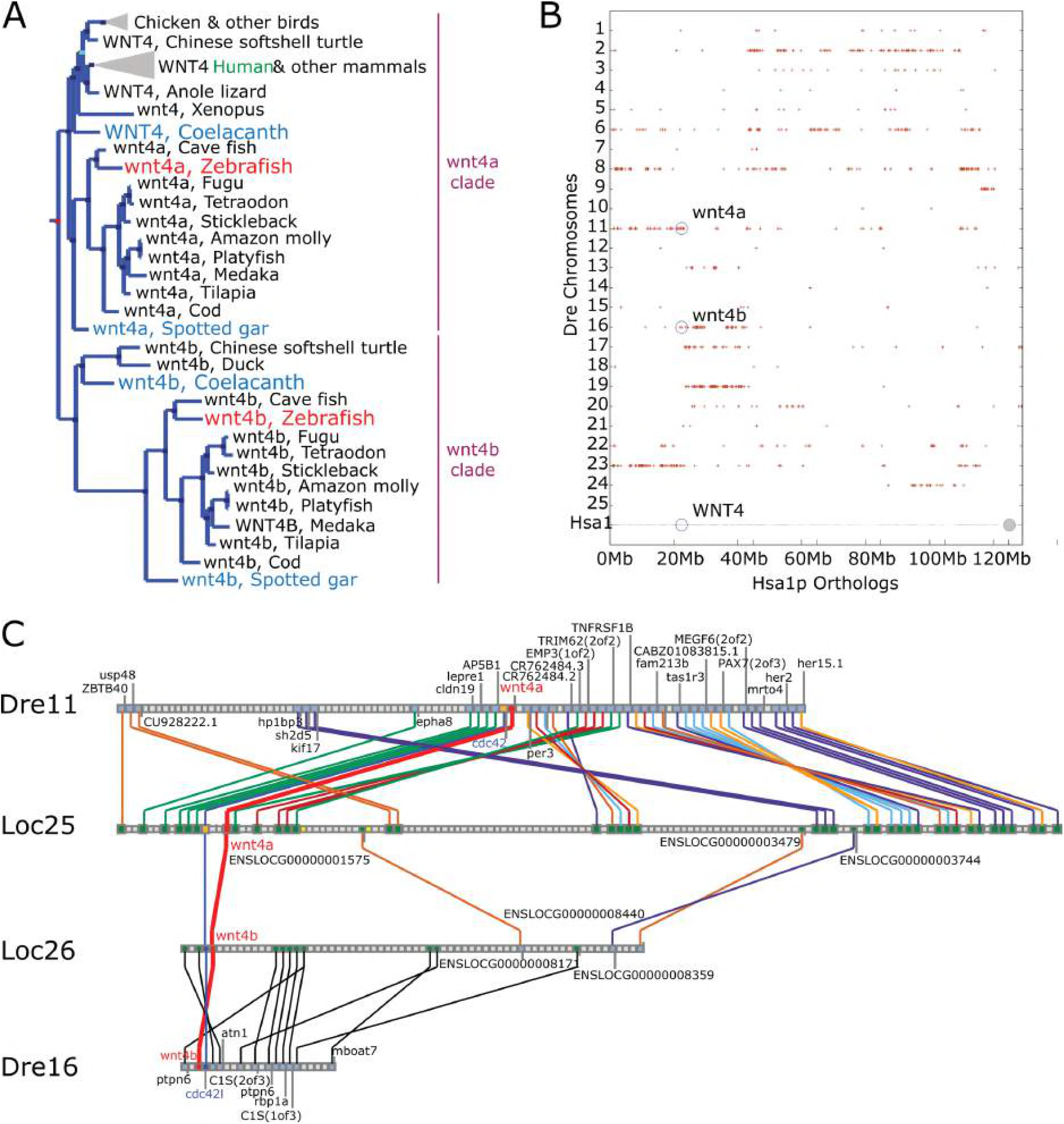
Teleost *wnt4a* is the ortholog of tetrapod WNT4. A. Phylogenetic analysis shows that vertebrates have two Wnt4-related clades (designated Wnt4a and Wnt4b). The Wnt4a clade includes teleosts and gar as ray-finned fish and coelacanth, birds, and mammals as lobe-finned fish. The Wnt4b clade also includes teleosts and gar as ray-finned fish as well as coelacanth and birds, but not mammals, as lobe-finned fish. This result shows that the duplication event that produced the *wnt4a* and *wnt4b* clades predated the divergence of ray-finned (e.g. gar, teleost) and lobed-finned (e.g. coelacanth, bird, mammal) lineages. B. A dot plot comparing zebrafish othologs and paralogs of genes on the short arm of human chromosome-1 (Hsa1p) shows conserved syntenies along zebrafish chromosome Dre11 (*wnt4a*) and Dre16 (*wnt4b*). C. Conserved synteny analysis shows that the zebrafish chromosome segments containing *wnt4a* and *wnt4b* are orthologous to regions on different spotted gar chromosomes, and that these two spotted gar chromosome show ancient paralogy. Based on phylogenetic and conserved synteny analyses, *wnt4a* and *wnt4b* were both in the last common ancestor of zebrafish and humans but mammals lost the ortholog of *wnt4b,* and *wnt4a* in teleosts is the ortholog of WNT4 in mammals.

Several lines of evidence argue that *wnt4a* and *wnt4b* have their origin in the two rounds of vertebrate genome duplication (VGD1 and VGD2), but not from retrotransposition, a simple one-gene duplication event or the teleost genome duplication event. First, the presence of introns in orthologous locations in both *wnt4* genes rules out the origin of either gene by retrotransposition. Second, analysis of conserved syntenies shows that *wnt4a* is located on zebrafish (*D. rerio*) chromosome Dre11 adjacent to *cdc42* while *wnt4b* is adjacent to *cdc42l* on Dre16, arguing that *wnt4a* and *wnt4b* arose from a genomic event more complicated than a simple one-gene tandem duplication (Amores *et al*. 1998) Figure 1B. Third, the teleost duplication event ohnolog of *wnt4a*-containing Dre11 is Dre23, while that of *wnt4b*-containing Dre16 is Dre19. (Amores *et al*. 1998) Figure 1B). Finally, the portion of Dre11 that contains *wnt4a* is orthologous to spotted gar (*Lepisosteus oculatus*) chromosome Loc25, which contains gar *wnt4a*, while the portion of Dre16 that contains *wnt4b* is orthologous to Loc26, which contains gar *wnt4b* (Figure 1C), as expected from whole genome duplication but not by tandem duplication. The finding that Loc25 and Loc26 are at least in part paralogous (Figure 1C) and that the gar lineage did not experience a genome duplication event after the divergence of ray-finned and lobe-finned vertebrates (Amores *et al*. 2011; Braasch *et al*. 2016) are as predicted by the hypothesis that *wnt4a* and *wnt4b* arose in one of the two genome duplication events at the base of the vertebrate radiation (Dehal and Boore 2005; Smith and Keinath 2015) and the *WNT4B* gene was lost in the mammalian lineage after it split from the bird lineage. We therefore conclude that the zebrafish *wnt4a* gene is the ortholog of the human gene *WNT4.*

### Early gonadal somatic cells express *wnt4a*

In mice, both XX and XY individuals express *Wnt4* in the early bipotential gonads (nine days post conception); thereafter, male gonads suppress *Wnt4* expression but female gonads maintain *Wnt4* expression (Vainio *et al*. 1999). We therefore used reverse transcription polymerase chain reaction (RT-PCR) to determine if the expression of either *wnt4a* or *wnt4b* were sexually dimorphic in adult and juvenile zebrafish gonads. Experiments detected *wnt4a* but not *wnt4b* in the ovary and *wnt4b* but not *wnt4a* in the testis in both adult and juvenile gonads (Supplemental Figure 1A, 1B). Thus, like mammalian *WNT4, wnt4a* in zebrafish appears to be associated with ovarian development.

We next asked where *wnt4a* is expressed in larval zebrafish gonads bracketing the sex determination and early sex differentiation period between 10 and 25dpf. Fluorescent RNA *in situ* hybridization experiments on wild-type gonads detected *wnt4a* expression in germ cells and in somatic gonad cells in all 17 individuals examined at 10dpf and 12dpf, though levels appeared to be higher in 10dpf gonads (Figures 2A, 2A’, n=8) relative to 12dpf wild-type gonads (Figures 2B, 2B’, n=9). In contrast, we were unable to detect *wnt4a* expression in any gonads in 14dpf individuals (Figure 2C, 2C’, n=6). At 20dpf, *wnt4a* expression was no longer detected in germ cells and appeared in only a subset of somatic cells (Figure 2D, 2D’, n=4). Somatic cell-specific expression of *wnt4a* increased from 20dpf until 23dpf when *wnt4a* was highly expressed in all gonads (n=10), specifically in the somatic cells surrounding larger oocytes indicating a presumptive ovary (Figure 2E, 2E’). Less expression was found surrounding smaller oocytes or cyst-like divisions of a presumptive testis (Figure 2F, 2F’). At 25dpf, when gonads had committed to the ovary or testis fate, which can be distinguished based on the numbers of oocytes present (Uchida *et al*. 2002), *wnt4a* was detected only in female gonads (Figure 2G, 2G’, n=6) and was no longer detected in developing male gonads (Figure 2H, 2H’, n=10). This sexually dimorphic expression of *wnt4a* continued throughout adulthood (Supplemental Figure 1A). We conclude that *wnt4a* is expressed in a dynamic, sex non-specific pattern in early gonads, and that by 25 dpf onward, its expression is limited to somatic cells in ovaries.

**Figure 2.**
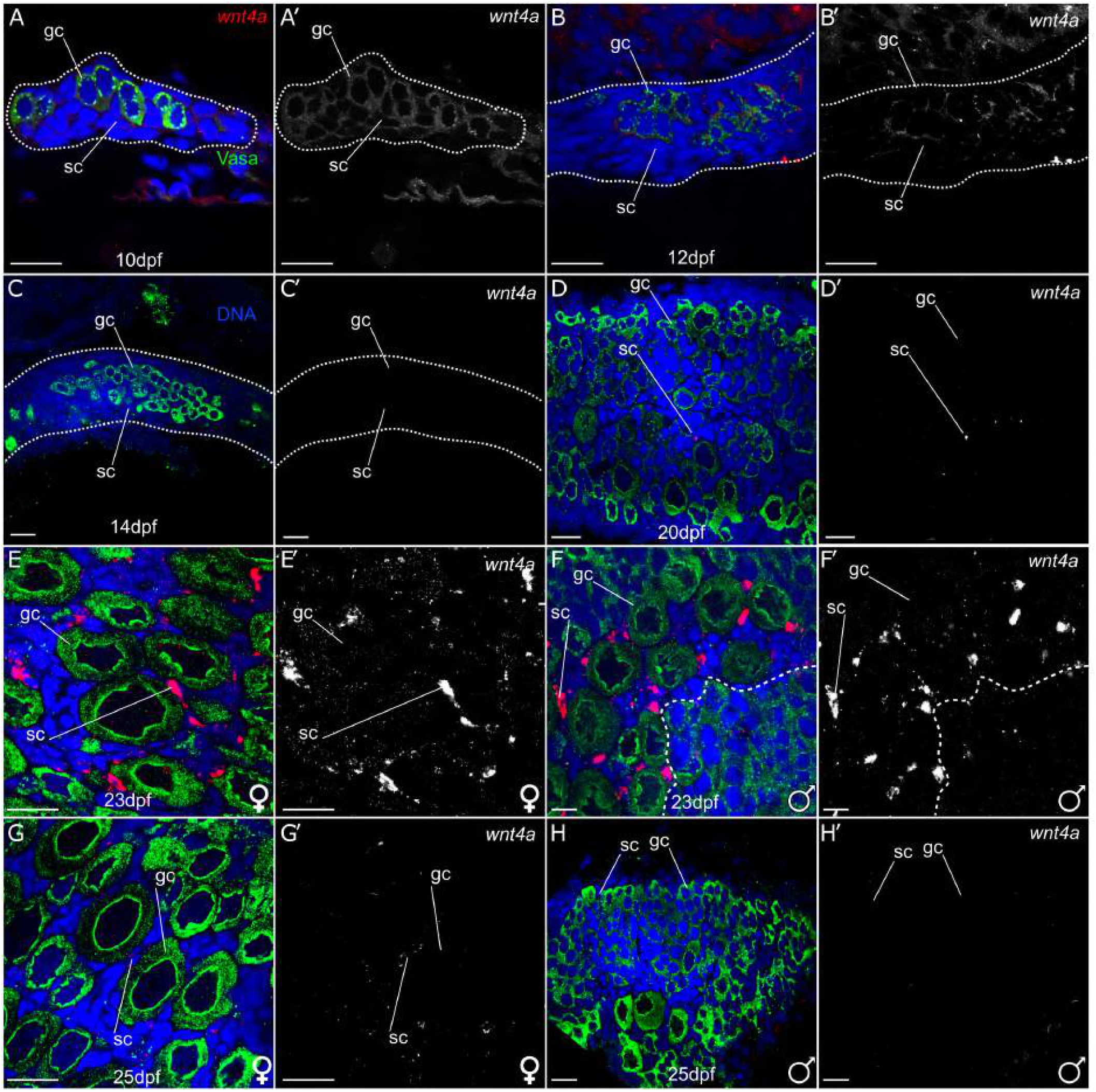
*wnt4a* is expressed in early zebrafish gonads. Confocal images of isolated gonads stained for *wnt4a* mRNA (red), Vasa to label germ cells (green), nuclei (blue), (A-H) or *wnt4a* RNA only (A’-H’), and gonadal tissue outlined by a dotted line (A-C). At 10dpf (A, A’, n=8) and 12dpf (B, B’, n=9) *wnt4a* RNA was detected ubiquitously in germ cells (gc) and somatic cells (sc). In contrast, *wnt4a* RNA was not detected in gonadal cells at 14dpf (C, C’, n=6). At 20dpf, *wnt4a* RNA was detected in a small subset of somatic cells (D, D’, n=4). At 23dpf (n=10), *wnt4a* expression was detected in somatic cells both in presumptive ovaries (E, E’) and in gonads that are transitioning to testes, with cyst-like divisions to the right of the dashed line (F, F’). At 25dpf, *wnt4a* mRNA was detected in the developing ovary (G, G’, n=6) but not in gonads that appeared to be transitioning to testes (H, H’, n=10). Scale bars = 20 µm.

We next asked if we could identify the somatic cell type that expresses *wnt4a* at 23dpf and 25dpf. At this stage, *wnt4a*-expressing cells associated closely with stage 1b oocytes (20µm – 140µm) and were therefore likely to be either theca cells or granulosa cells. To distinguish between these possibilities, we used the transgenic reporter line Tg(*cyp19a1a:egfp*), which expresses GFP in theca cells that surround stage 1b oocytes (Dranow *et al*. 2016). Results showed that *wnt4a* expressing cells did not also express GFP (n=3, Supplemental Figure 2). We conclude that *wnt4a* is not expressed in theca cells, but rather in another component of the gonadal soma, likely granulosa cells, although another gonadal cell type cannot be excluded.

### *wnt4a* mutants develop predominantly as males

Results so far indicated that *wnt4a* is predominantly expressed in female somatic gonad cells. This is consistent with the hypothesis that Wnt4a plays a role in female sex determination in zebrafish. To test this possibility, we analyzed the phenotype of two ENU-induced alleles of *wnt4a*: *wnt4a(fh294)* and *wnt4a(fh295)*, which were identified by TILLING (Moens 2009; Choe *et al*. 2013). The *wnt4(fh294)* and *wnt4a(fh295)* alleles are nonsense point mutations that are predicted to truncate the C-terminus of Wnt4a protein (Figure 3A). The *wnt4a(fh294)* and *wnt4a(fh295)* mutations result in the deletion of ten or one of the conserved cysteines, respectively, that are present in the carboxy-terminus of the Wnt4a protein, and that are necessary for proper folding of WNT proteins (Miller 2002). Without these residues, the binding of Wnts to the Frizzled receptor is likely to be disrupted (Janda *et al*. 2012). Importantly, deletion of the carboxy-terminal half of the Xenopus *Xwnt*-8 gene results in a partial protein that has dominant-negative, cell non-autonomous activity, perhaps because it interferes with productive interations between the wild-type XWnt8 ligand and its receptor (Hoppler *et al*. 1996); given the high sequence conservation of Wnt ligands, it was therefore possible that carboxy-terminal deletions of Wnt4a may behave similarly. To investigate this possibility, we used CRISPR/Cas9 to generate additional mutations targeted to the N-terminus. The *wnt4a(uc*55) and *wnt4a(uc56)* alleles resulted from a 17bp and 23bp insertion in exon two, respectively, and are therefore predicted to cause translational frame shifts, (Figure 3A, Supplemental Figure 3) and hence to be strong loss-of-function alleles. In support of this prediction, we could not detect any wild-type *wnt4a* mRNA by reverse transcription-polymerase chain reaction analysis (RT-PCR) in *wnt4a(uc55)* mutants, suggesting that the mutant transcript is subject to nonsense mediated decay (data not shown), suggesting that the mutant transcript is subject to nonsense mediated decay.

**Figure 3.**
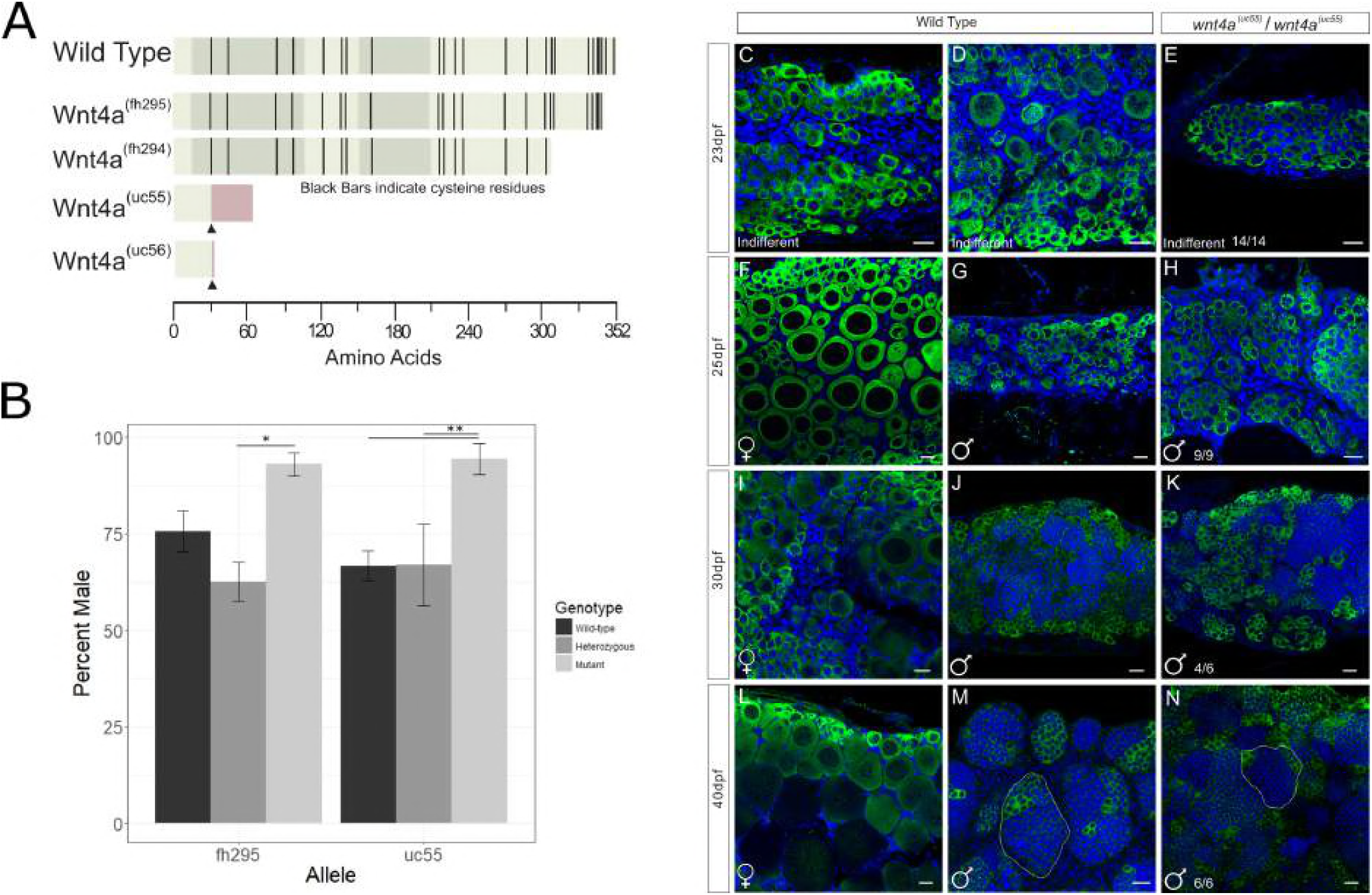
Mutant *wnt4a* alleles result in male-biased populations. A. Wnt4a is a 352 amino acid protein with five exons, indicated by the alternating shaded regions. Protein structures predicted to arise from each allele are indicated by truncation, *wnt4a(fh295/fh294)* (ENU induced mutation) or insertion (CRISPR-arrowhead) resulting in missense protein coding (red bar) in uc55/uc56. B. Sex ratios in populations of homozygous *wnt4a(fh295)* and *wnt4a(uc55)* mutants were significantly male biased by anova (* = p<0.05, ** = p<0.01, n=3 replicates). Comparison of representative wild-type (C, D, F, G, I, J, L, M) and *wnt4a* mutant (E, H, K, N) gonads stained for Vasa, to identifiy germ cells (green), and DAPI, to label nuclei (blue), at various ages post-fertilization (dpf). At 23dpf, *wnt4a* mutants (E) and wild-types (C, D) both have indifferent gonad morphology. In contrast, the majority of *wnt4a* mutant gonads from 25dpf animals and older (H, K, N) had a morphology that is indistinguishable of wild-type testes (G, J, M), but not wild-type ovaries (F, I, L). Scale bars = 20µm.

We first asked if the CRISPR-induced *wnt4a* mutants were viable. We crossed parents that were heterozygous for each mutant allele, genotyped the resulting offspring at three months of age, and determined their phenotypic sex. For all four alleles, we found the expected Mendellian1:2:1 ratio of the three possible genotypes (*wnt4a*^+/+^: *wnt4a*^+/-^: *wnt4a*^-/-^; Supplemental Table 2; Chi-Squared test). In contrast to mammals, where *Wnt4* mutants are embryonic lethal, *wnt4a* loss-of-function zebrafish mutants are viable (Supplemental Figure 4). We next determined if loss of Wnt4a function affected sex ratios (Figure 3B). In the *wnt4(fh295)* in-cross population, *wnt4(fh295)* wild-type fish were 75.7% male, *wnt4(fh295)* heterozygous fish were 62.6% male whereas the *wnt4(fh295)* homozygous mutant fish were 93% male (n=3, p<0.05, anova). Similarly, the *wnt4a(uc55)* mutation resulted in similar ratios (wild-type, heterozygous and homozygous mutant fish were 66.6%, 76% and 94% male, respectively, n=3, p<0.01, anova). Homozygous mutants for *wnt4a(fh294)* and *wnt4a(uc56)* also had male sex bias (98.6% n=141/143 and 100% male n=78/78, respectively). Finally, *wnt4a(uc55)/wnt4a(fh295)* trans-heterozygous fish had a similar male bias compared to homozygous single mutants (96% male; n= 22/23). These data support two main conclusions. First, these results indicate that Wnt4a promotes ovary development but is not absolutely required for ovary development. Second, because the *wnt4a(fh295)* mutants had the same magnitude of effect on sex ratios as the loss-of-function allele *wnt4(uc55)*, we conclude that the ENU induced alleles, *wnt4a(fh294)* and *wnt4a(fh295),* are also loss-of-function.

### Wnt4a is involved in primary sex determination and/or differentiation

In mammals, WNT4 is required during female primary sex determination (Vainio *et al*. 1999). In zebrafish, it is not known with certainty when definitive primary sex determination occurs, but it likely occurs prior to 20dpf, because this is the time at which oocytes present in the bipotential gonad begin to die by apoptosis in presumptive males (Takahashi 1977; Uchida *et al*. 2002). The hypothesis that *wnt4a* is required for primary sex determination and/or differentiation in zebrafish predicts that oocyte apoptosis will initiate in the majority of mutants at about the same time as it does in wild-type males, but in a greater proportion of the population. Alternatively, the hypothesis that *wnt4a* is instead required to maintain female sex differentiation predicts that many animals should begin to develop as females, but then revert to male phenotype during the early juvenile stage, as occurs in *bmp15* mutants (Dranow *et al*. 2016). We therefore compared gonad development between wild-type and *wnt4a* mutants between 23-40dpf (Figure 3C-3N). Results showed that the majority of *wnt4a* mutant gonads were morphologically similar to wild-type males at all stages analyzed (compare Figure panel 3G, 3J, 3M to panel 3H, 3K, 3N), but not to wild-type females (compare Figure panel 3F, 3I, 3L to panel 3H, 3K, 3N). In particular, early stage oocytes were present in all gonads at 23dpf regardless of genotype, but by 25dpf, mutant gonads appeared to contain predominantly pre-meiotic germ cells, which have nuclei containing a single large nucleolous, similar to those found in presumed wild-type males (Figure 3H). By 40dpf, all mutant gonads had a morphology that was indistinguishable from a wild-type testis, where germ cells are organized into tubules (compare Figure 3M to 3N). These data thus argue that Wnt4a is involved in primary sex determination and/or differentiation rather than in the maintenance of a female phenotype.

### Wnt4a mutants are unable to release gametes

The ovaries and testes of *wnt4a* mutant adults are morphologically indistinguishable from those of their wild-type siblings (Supplemental Figure 5). It was therefore that surprising that neither mutant males nor mutant females produced progeny when mated to each other or to wild-type fish. For example, *wnt4a(*uc55) mutant males stimulated wild-type females to lay eggs, but no eggs were fertilized (n=155 eggs) and for *wnt4a(fh294)*, nine homozygous wild-type male siblings and nine homozygous mutant males were individually crossed to 2-3 AB wild-type females. We found that mating with wild-type males produce 349/504 (69.2%) viable offspring, while those with mutant males produced only unfertilized eggs (n=471).

Because our histological analysis showed that mature sperm were present in the testes of mutant males (Supplemental Figure 5B), we next attempted to expel sperm from mutants by gentle squeezing. We found that *wnt4a(uc55)* and *wnt4a(fh294)* wild-type control males released sperm (n=8/9 and 16/21, respectively), but that mutant males did not (n=0/10 and 0/16 respectively). Finally, we used an *in vitro* fertilization assay to compare fertilization rates of mutant and wild-type sperm isolated from dissected and macerated testes. We found that, consistent with our histological analysis, dissection-isolated mutant sperm had similar fertilization rates to sperm isolated from heterozygous or homozygous wild-type males (for *wnt4a(uc55)*: 56.1±35.6% for mutants vs. 79.3±7.2%, for wild-types; P=0.46, Student’s T-test and for *wnt4a(fh294)*: 72.7% for mutant males and 52.2% for wild types, P=0.71.)

Similarly, histological analysis showed that ovaries in the mutant females obtained contained all stages of oocytes, including mature eggs (Supplemental Figure 5D), yet *wnt4a(uc55)* mutant females failed to release eggs when mated to wild-type males (0/5 mating pairs). In contrast, two of three heterozygous control females released eggs when mated to a wild-type male. We next tested if we could recover eggs by gentle squeezing and found that, although two of three control females released eggs, no *wnt4a(uc55)* mutant females released eggs (n=0/5). Finally, we tested if we could recover mature eggs from dissected ovaries. We found that eggs dissected from *wnt4a(uc55)* mutant females yielded viable zygotes when fertilized by wild-type sperm, though at a lower rate than those isolated from heterozygous females (16.4 ± 6.2%, n=3 mutants females vs. 61.4 ± 38.1%, n=4 heterozygotes females, P=0.41 Student T-test). These results show that the infertility of *wnt4a* mutant males and females is not due to a defect in gametogenesis, and we hypothesized that it was due instead to an inability of mutants to release their gametes.

### Male and female infertility is caused by reproductive duct malformation

Given that *wnt4a* mutant zebrafish cannot expel their gametes, we asked if they had defects in the formation of the reproductive ducts. In wild-type males, each testis connects to the genital orifice by the duct deferens (DD), which extends posteriorly from the testis and fuses with the genital orifice to form the fused duct deferens (FDD, Figure 4A, 4B). We analyzed duct formation first by histology using Hematoxylin and Eosin (H&E) stained paraffin sections. We found that although the duct deferens initiated development in mutant males, their extension was variable and the ducts failed to fuse (n=3; Figure 4A’, 4B’).

**Figure 4.**
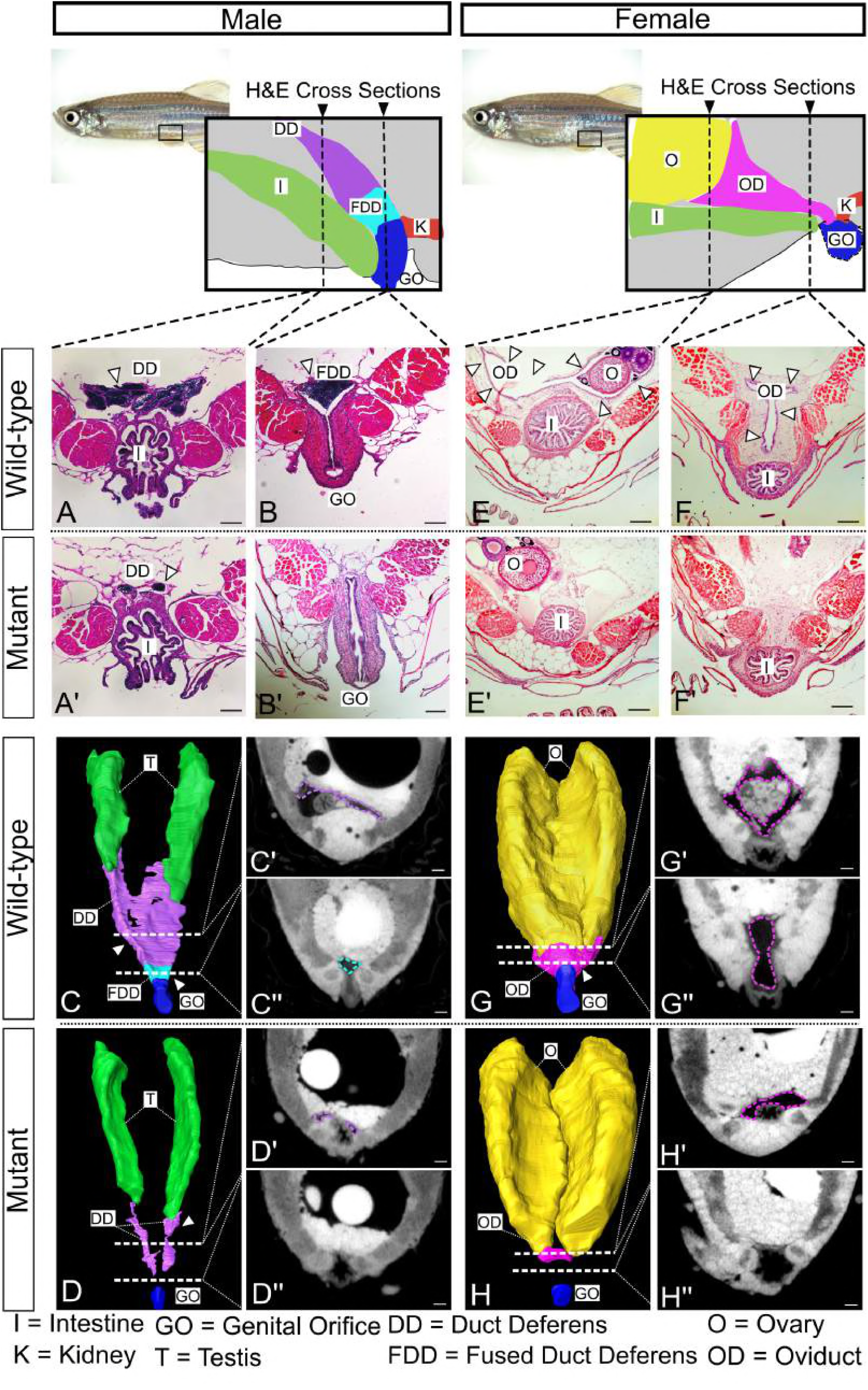
Duct morphology in *wnt4a* mutant and wild-type males. From the posterior end of each testis, wild-type males developed duct deferens (A) that joined to form the fused duct deferens (B). Mutant males, however, failed to form a full duct deferens (A’) or a fused duct deferens (B’). 3D renderings built from individual traces of sections (C’, C’’ etc) of the wild-type ducts (C) and mutant ducts (D) show that the mutant ducts never fully connected to the genital orifice. H&E stained sections of wild-type females showed an oviduct that wrapped around the posterior of the ovaries (E, G, G’) and extended ventroposteriorly (F, G, G’’) out to the genital orifice. Mutant females, however, failed to organize an oviduct around the ovary (E, H, H’) and did not form a connective duct to the genital orifice (F, H, H’’). Scale bar = 100µm (See movies in Supplemental Figures 6 and 7.)

To increase the resolution of assessing duct development in mutants, we next utilized micro Computed Tomography (microCT) to render 3D representations of the reproductive ducts in both wild-type and mutant adults. For this analysis, we traced the structures of interest in individual slices (Figure 4C’, 4C’’) to build a 3D model of the complete duct structure (Figure 4C). In wild-type animals, as expected, we could identify all parts of the reproductive ducts (Figure 4C, 4C’, 4C’’). By contrast, in *wnt4a(uc55)* mutant males, we could identified DD but no duct fusion or connection to the genital orifice (Figure 4D, 4D’; Supplemental Figure 6).

In females, histology and microCT analysis revealed a similar defect in reproductive duct development in *wnt4a* mutants. In wild-type females, the oviduct wrapped around the posterior end of the ovary and extended ventroposteriorly until it connected to the genital orifice (Figure 4E, 4F, 4G). In mutant females, however, the small amount of oviduct tissue present did not fully envelop the posterior end of the ovaries (Figures 4E’, 4H, 4H’; Supplemental Figure 7), and failed to extend towards the genital orifice (Figure 4F’, 4H, 4H’’; Supplemental Figure 7). Together, these analyses explain mutant sterility and show that Wnt4a is required not for the specification of reproductive duct development, but is likely required for the growth and/or extension of the reproductive duct primordium in both male and female zebrafish.

To further understand how Wnt4a regulates duct development, we first determined when the ducts form during larval development. We scored duct formation in wild-type males using serial H&E sections. We evaluated males at four ages from 25dpf to 55dpf and discovered that the ducts appear to originate at the posterior end of the testis before 25dpf and elongate towards the genital orifice. Reproductive ducts of wild-type males had reached and fused to the genital orifice by 55dpf (Supplemental Figure 8).

Having established the schedule of duct development, we wanted to learn in which tissues *wnt4a* acts to cause duct elongation: Does Wnt4a act in the extending duct? Or in the space through which it grows? Or in the target at the vent? To find out, we analyzed serial transverse sections of wild-type males starting anterior to the gonad and ending posterior to the genital orifice with alternate sections taken for H&E histology (see Supplementary Figure 8) and expression analysis of *wnt4a* by in situ hybridization. At 25 and 35dpf, *wnt4a* expression appeared not in the extending duct, but around the vent and developing genital orifice, (Figure 5B, 5D). While no *wnt4a* expression was detected in cells of the DD primordium at 25dpf (Figure 5A), by 35dpf *wnt4a* expression appeared in cells located ventral to the developing DD (Figure 5C). Importantly, the domain of *wnt4a* expression preceded the arrival of the ducts to this region.

**Figure 5.**
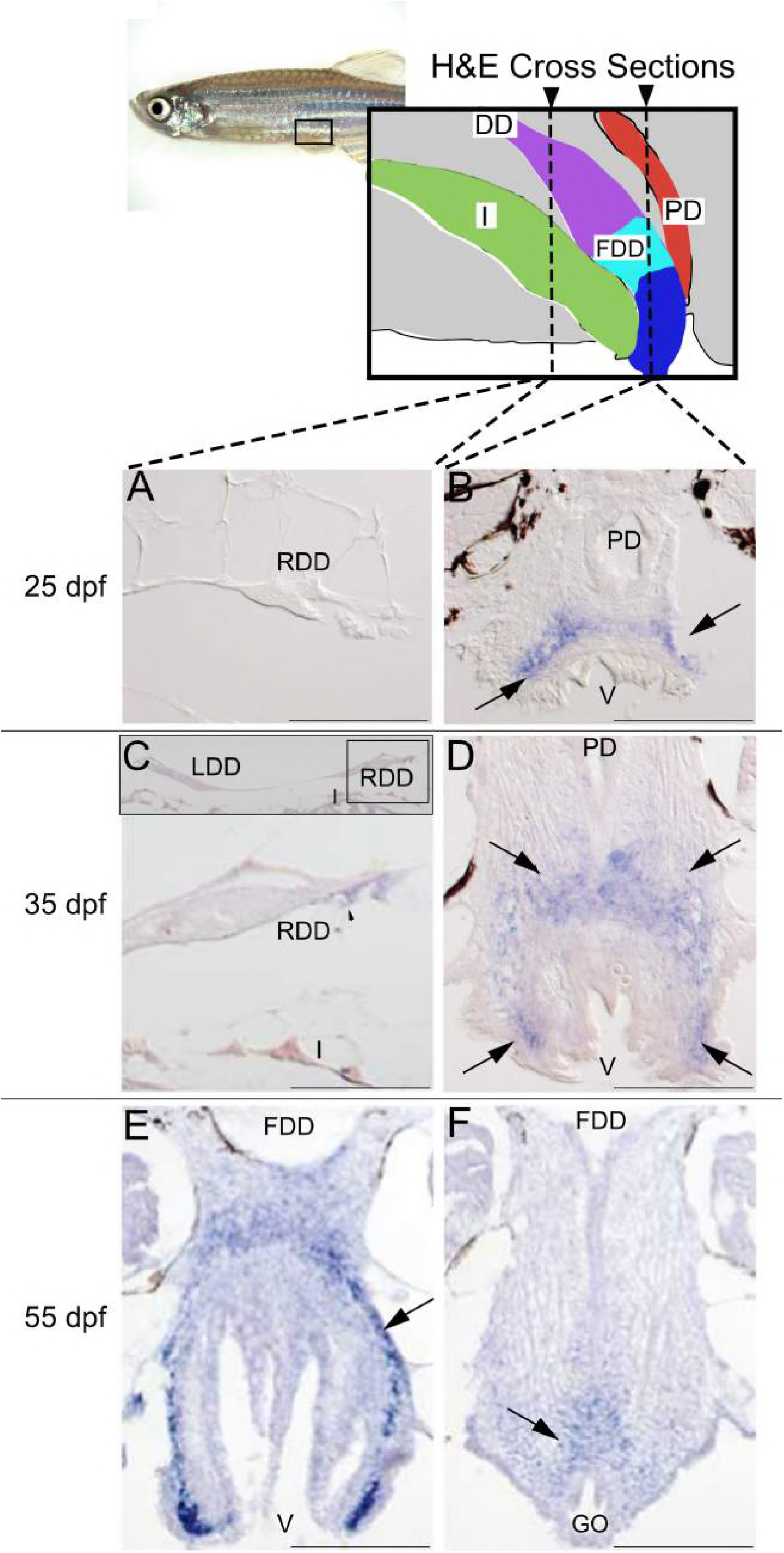
Expression of *wnt4a* during reproductive duct development in zebrafish wild-type AB strain males. Alternate serial cross sections of wild-type males were stained by H&E to follow duct growth (see Supplementary Figure 8) or prepared for *in situ* hybridization to reveal *wnt4a* expression, shown here. A. Expression analysis showed that at 25dpf, the duct deferens (cross section of the right duct shown here) lacked *wnt4a* expression. B. At 25dpf, however, *wnt4a* expression (black arrows) appeared dorsal to the posterior vent, just anterior to the eventual connection of the fused duct deferens and the genital orifice. C. At 35dpf, the duct deferens continued to show little *wnt4a* expression. (Insert shows the left and right duct deferens dorsal to the intestine.) D. At 35dpf, *wnt4a* expression (black arrows) appeared dorsal and lateral to the posterior vent, just anterior to the eventual connection of the fused ducti deferens and the genital orifice. E. At 55dpf in a section just anterior to the fusion of the duct deferens to the vent, *wnt4a* expression (black arrow) appeared dorsal to the posterior vent, just anterior to the connection of the fused ducti deferens and genital orifice. F. At 55dpf in a section at the level of the connection of the fused duct and the vent, *wnt4a* expression (black arrow) appeared just dorsal to the genital orifice. (PD = pronephric duct, FDD = fused ducti deferens, GO = genital orifice, DD = ducti deferens, RDD = right duct deferens, LDD = left duct deferens, I = intestine, V = vent) Scale bar = 100µm.

Serial sections showed that wild-type DD connected to the genital orifice between 45 and 55 dpf (Supplementary Figure 8). Concurrent analysis of alternating sections showed *wnt4a* expression persisted in tissue surrounding the the vent and genital orifice (Figure 5E, 5F). We conclude that in wild-type male zebrafish, *wnt4a* expression occurs in the developing genital orifice but not in the extending DD, raising the hypothesis that Wnt4a might act as a diffusible signal that encourages DD outgrowth.

To further characterize the role of Wnt4a in duct development, we examined duct elongation in *wnt4a* mutants over time. Results showed that the DD elongated more slowly in *wnt4a* mutant males than wild-type. The FDD did not connect the DD to the genital orifice by 55dpf nor was a connection found in elderly 2-year old fish (Supplemental Figure 8). These results suggest that *wnt4a* expression at the genital orifice is essential for reproductive duct growth and/or elongation and for formation of the FDD. The failure of ducts to connect to the genital orifice explains our earlier observations that *wnt4a* mutants are sterile despite their ability to make fertile eggs and sperm. The *wnt4a* expression domain anterior to the eventual connection of the FDD and the genital orifice supports the hypothesis that tissues around the genital orifice likely secretes Wnt4a protein, and thus signal duct growth, elongation and connection to the exterior. Thus, as in mammals, Wnt4a may coodinate directional cell migration and extention of the reproductive duct (Prunskaite-Hyyryläinen *et al*. 2015).

## DISCUSSION

After more than four decades of use as a major model organism, the mechanism of sex determination in domesticated zebrafish is still not clear. While a major sex chromosome has been identified in wild zebrafish, this sex-determining locus appears to have been lost during the domestication of zebrafish strains that are widely used in the laboratory (Wilson *et al*. 2014). Regardless, there is mounting evidence that many, if not most, genes that play key roles in sex determination in mammals animals play similar roles in zebrafish. As an example, the double-sex and mab3 related transcription factor Dmrt1, a highly conserved regulator of male development across metazoans, is required for normal male development in zebrafish (Lin *et al*. 2017; Webster *et al.* 2017). Similarly, in vertebrates, WNT4 signaling plays a key role in female sex determination and accumulating evidence argues that canonical Wnt signaling is also required for female sex determination in zebrafish, though the specific Wnt ligand had not been previously identified (Zhang *et al*. 2011; Sreenivasan *et al*. 2014). Experiments reported here show that the zebrafish ortholog of mammalian WNT4, *wnt4a*, is required for normal female sex ratios, strongly suggesting that it plays a role in female sex determination but is not required for female sex determination because a small percentage of *wnt4a* mutants develop as females. In addition, while WNT4 in mammals is required for the development of reproductive ducts in the female, but not the male (Vainio *et al*. 1999), we have shown here that in zebrafish, Wnt4a is required for reproductive duct development in both females and males. Together, these results provide further evidence that the underlying mechanisms for sex determination and/or differentiation are well conserved between teleosts and tetrapods.

The zebrafish genome contains two *Wnt4*-related genes, *wnt4a* and *wnt4b*, while the mammalian genome contains a single *WNT4* gene (Ungar *et al*. 1995; Liu *et al*. 2000). Although many gene duplicates in teleosts are the result of an additional whole genome duplication event that occurred after the teleost and tetrapod lineages diverged (Amores *et al*. 1998; Postlethwait *et al*. 1998; Jaillon *et al*. 2004), our phylogenetic analysis argues that the duplication event that produced *wnt4a* and *wnt4b* predated the teleost-tetrapod divergence. Specifically, while mammals have only a single copy of Wnt4, coelacanth and birds (among basally diverging lobe-finned fish) and spotted gar (among basally diverging ray-finned fish) contain two orthologs of Wnt4. Based on sequence comparisons and analysis of conserved syntenies, it is clear that the single *Wnt4* copy that remains in mammals is the ortholog of the teleost *wnt4a* gene, indicating that the ortholog of *wnt4b* was lost at some point after the mammalian linage diverged from the turtle and bird lineages. Thus, although we do not propose a name change for practical reasons, in principle, the human gene should be called *WNT4A* to match its teleost ortholog, or the teleost gene should be called simply *wnt4* to match its mammalian ortholog.

The early gonad in mammals is bipotential, and expresses *Wnt4* initially in the mesonephros underlying the *Fgf9*-expressing gonadal epithelium (Vainio *et al*. 1999). Mutational analysis has shown that Wnt4 and Fgf9 are mutually antagonistic during mammalian sex determination: loss of *Wnt4* function in XX mammals leads to upregulation of *Fgf9* and partial female-to-male sex reversal (Vainio *et al*. 1999), whereas loss of *Fgf9* in XY individuals results in stabilized expression of *Wnt4* and partial male-to-female sex reversal (Kim *et al*. 2006). During normal development, *Sox9* expression in the gonad, which is initiated by the mammalian Y-linked male sex-determinant SRY, leads to up-regulation of *Fgf9*, which in turn down-regulates *Wnt4*. In contrast, in the absence of *Sox9* expression, as occurs normally in XX mammals, *Wnt4* represses *Fgf9*, thus promoting female development (Kim *et al*. 2006).

We have shown here that the phenotypes caused by *wnt4a* mutations in zebrafish, such as masculinization of the gonad and disturbed sex duct development, parallel those of *Wnt4* mutant mammals, yet it is not clear whether the mechanisms by which Wnt4a promotes ovarian and gonadal duct development are conserved. Our results clearly show that, as in mammals, *wnt4a* is expressed in somatic gonadal cells during the bipotential phase of gonad development. However, while the genome of a basal teleost, the spotted gar, contains an ortholog of Fgf9 (Braasch *et al*. 2016), orthologs of Fgf9 have not been found in the genomes of other teleosts, included zebrafish (Itoh and Konishi 2007), suggesting that this gene was lost during early teleost evolution. It therefore remains to be determined if another Fgf ligand plays a similar role in teleosts to that of mammalian Fgf9 in opposing the action of Wnt4a during sex determination.

Three noteworthy features differ between the phenotypes of zebrafish and mammalian Wnt4 mutants. First, loss-of-function *Wnt4* mutants in mice and humans are lethal (Vainio *et al*. 1999; Mandel *et al*. 2008), whereas zebrafish *wnt4a* mutants are viable. It is likely that in mammals the lethal phenotype of *Wnt4* mutants is due to an additional and essential function of Wnt4 during the development of non-gonadal tissues. If so, then the viability of *wnt4a* mutant zebrafish may be the result of Wnt4b function in non-gonadal tissue development. For example, in mouse and zebrafish, Wnt4 and *wnt4b*, respectively, are expressed in the floorplate of the spinal cord and brain (Parr *et al*. 1993; Liu *et al*. 2000; Agalliu *et al*. 2009; Duncan *et al*. 2015). In addition, lethality of WNT4 mutant mice and humans is likely due to kidney failure (Vainio *et al*. 1999; Mandel *et al*. 2008). To date, however, the expression of either *wnt4a* or *wnt4b* has not been reported in the pronephros, the zebrafish equivalent to the mammalian kidney. Regardless, it remains to be determined if simultaneous loss *wnt4a* and *wnt4b* in zebrafish will cause embryonic lethality.

Second, in mammals, XX Wnt4 mutants are partially sex reversed to males and germ cells undergo apoptosis, whereas in zebrafish, all *wnt4a* mutants produce functional gametes, including the 4-6% of Wnt4a mutants that develop as females. It is likely that this difference results from the observation that in mammals, gametes do not survive if the gonadal sex is opposite of the somatic sex, regardless of the direction of sex reversal (Uhlenhaut *et al*. 2009; Matson *et al*. 2011). In contrast, ample evidence shows that in many teleost, including zebrafish and medaka, the gamete type produced by premeiotic germ cells can readily switch to match the sexual phenotype of the somatic gonad, regardless of whether the phenotype matches the genetic sex of the animal (Yamamoto 1958; Dranow *et al*. 2013, 2016; Wong and Collodi 2013).

Third, unlike mammals, Wnt4a in zebrafish appears to facilitate, but is not essential for, female development, because a small percentage of *wnt4a* mutants develop normal ovaries. Two models could explain this difference. First, it is possible that Wnt4b can partial compensate for loss of Wnt4a during female sex determination or differentiation. Alternatively, it is possible that female development of *wnt4a* mutants is related to the numbers of oocytes that these individuals possess during the critical sex-determining window. During this time period (10dpf-20dpf), all zebrafish juveniles produce several early stage oocytes and mounting evidence shows that the number of oocytes an individual produces during the bipotential phase correlates with the eventual sex of the animal: animals that produce few or no oocytes become male, whereas those that produce many oocytes can become female (Uchida *et al*. 2002; Rodríguez-Marí *et al*. 2010; Dai *et al*. 2015). While it is not known for certain, it is likely that oocytes produce a signal that acts on the somatic gonad to promote female sex determination and absent a threshold amount, animals develop as males (Figure 6A). We therefore propose two general models that the role of Wnt4a during normal sex determination. First, it is possible that Wnt4a may regulate the sensitivity of the somatic gonad to the oocyte-produced signal, such that, in wild-type gonad, fewer oocytes are required to reach the critical threshold necessary to stabilize female sex determination relative to *wnt4a* mutant gonads (Figure 6B). Alternatively, Wnt4a may act on germ cells to regulate the level of signal produced (Figure 6C), either by directly regulating the amount of signal each oocyte produces or effecting the number of oocytes produced per animal. Regardless, our observation that some *wnt4a* mutants can develop as females suggests that above a certain level of signal, Wnt4a function is not required for female development.

**Figure 6.**
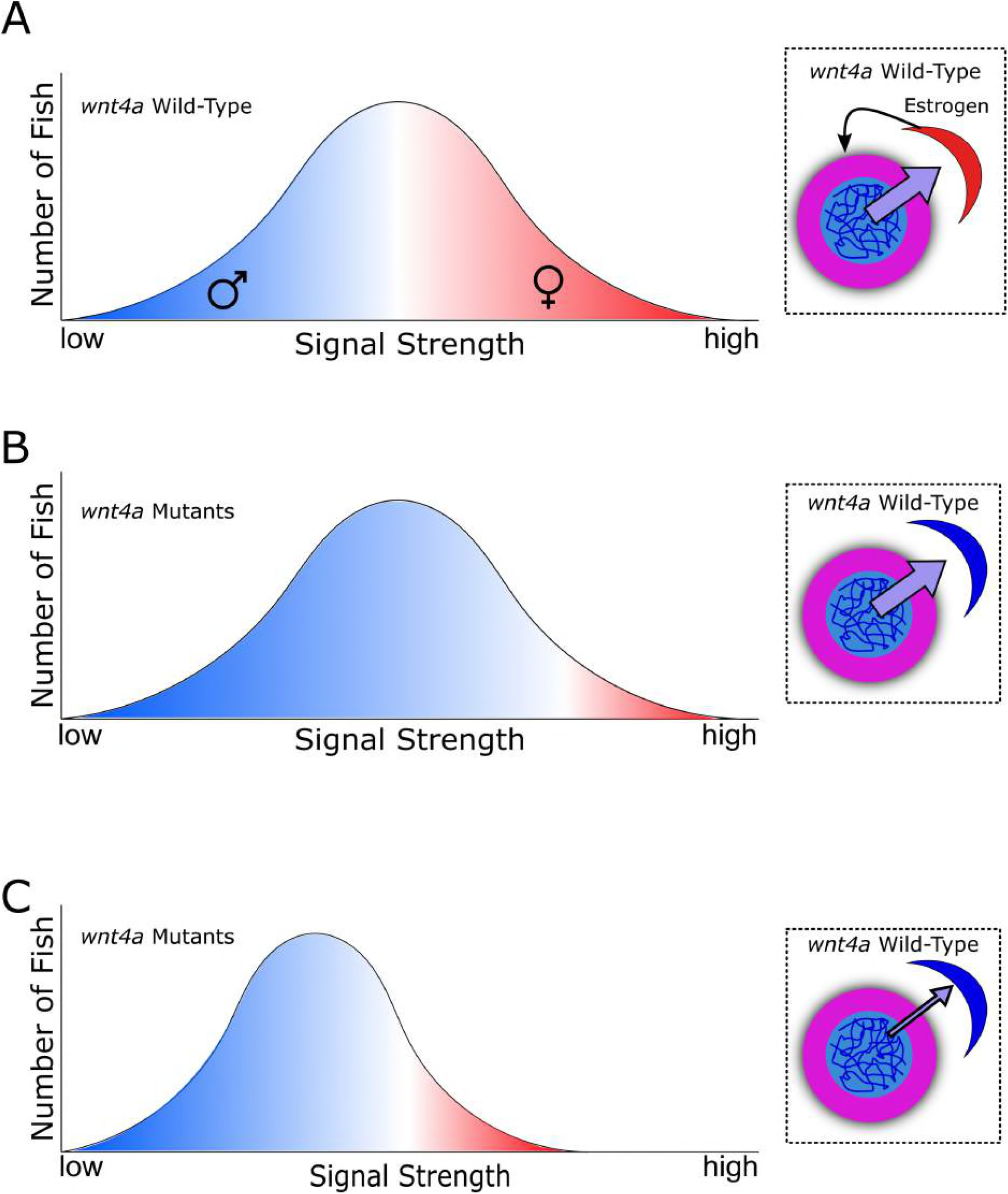
Models for how Wnt4a functions to promote female development. A. In wild-type animals, high concentrations of a signal produced by early-stage oocytes during the bipotential gonad stage (purple arrow) likely cause the gonadal soma to maintain production of estrogen (black arrow), which inhibits oocyte death and drives female sex determination. If the oocyte signal is too low, a male develops; if the signal exceeds a threshold, a female develops. In a wild-type population, this threshold and signal gradient produces about half males and half females. In A-C, the X-axis depicts the strength of the signal while the Y-axis plots the numbers of animals that produce a certain amount of signal. For simplicity, signal strength vs. fish number is assumed to follow a normal distribution. Color intensity reflects the probability an individual develops as a male (blue) or female (red). B. In this model, lack of Wnt4a desensitizes somatic gonad cells to the female-promoting oocyte signal, thereby raising the female-development threshold such that only those few *wnt4a* mutant animals that produce the highest signal (perhaps stochastically) can develop as females allowing most to become males. C. Alternatively, lack of Wnt4a causes oocytes to decrease the amount of female-promoting signal that they produce, such that fewer *wnt4a* mutants achieve the level required to sustain female development. Insets in B and C are graphical representations of the two models (oocyte in pink, somatic gonad cell in red or blue).

Finally, we have shown that both male and female *wnt4a* mutants are unable to release their gametes due to defects in reproductive duct development. In mammals, the reproductive ducts of males and females develop from separate embryonic structures, the Müllerian duct in females and the Wolffian duct in males. These reproductive duct anlagen initially develop in both males and females during early embryogenesis, but after definitive sex determination has occurred, the Müllerian ducts degenerate in males, while the Wolffian ducts degenerate in females. Loss of Wnt4 function in mice inhibits Müllerian duct formation in both males and females, but does not affect the development of the Wolffian ducts (Vainio *et al*. 1999). This finding suggests that the reproductive ducts in both male and female zebrafish are developmentally similar to the Müllerian ducts in mammals and may therefore share a common evolutionary origin. Recent studies in mice have shown that WNT4 regulates the direction of Müllerian duct precursor cell migration (Prunskaite-Hyyryläinen *et al.* 2015). How Wnt4a regulates ductal development in zebrafish remains to be determined.

In conclusion, results presented here establish that Wnt4 is likely a regulator of female sex determination and reproductive duct development in the last common ancestor of humans and zebrafish 450 million years ago. As such, these results provide further evidence that the core pathway for sex determination and differentiation in tetrapod vertebrates appears to be largely conserved in the teleost lineage.

## ACKNOWLEGDEMENTS

This work was made possible by generous funding from the following sources, T32ES0070599 (Michelle Kossack), T32HD008348 (Samantha High), R01 GM085318 (John Postlethwait), and IOS-1456737 (Bruce Draper). The Olympus FV1000 confocal used in this study was purchased using NIH Shared Instrumentation Grant 1S10RR019266-01. We thank the MCB Light Microscopy Imaging Facility, which is a UC-Davis Campus Core Research Facility, for the use of this microscope. MicroCT was supported by a pilot grant from the University of California-Davis Center for Molecular and Genomic Imaging. We would also like to thank the members of the Draper and Postlethwait labs especially Matt McFaul, Anastasia Utkina, Becky Wong, and Trevor Enright for their contributions.

## SUPPLEMENTAL TABLE LEGENDS

**Supplemental Table 1.**
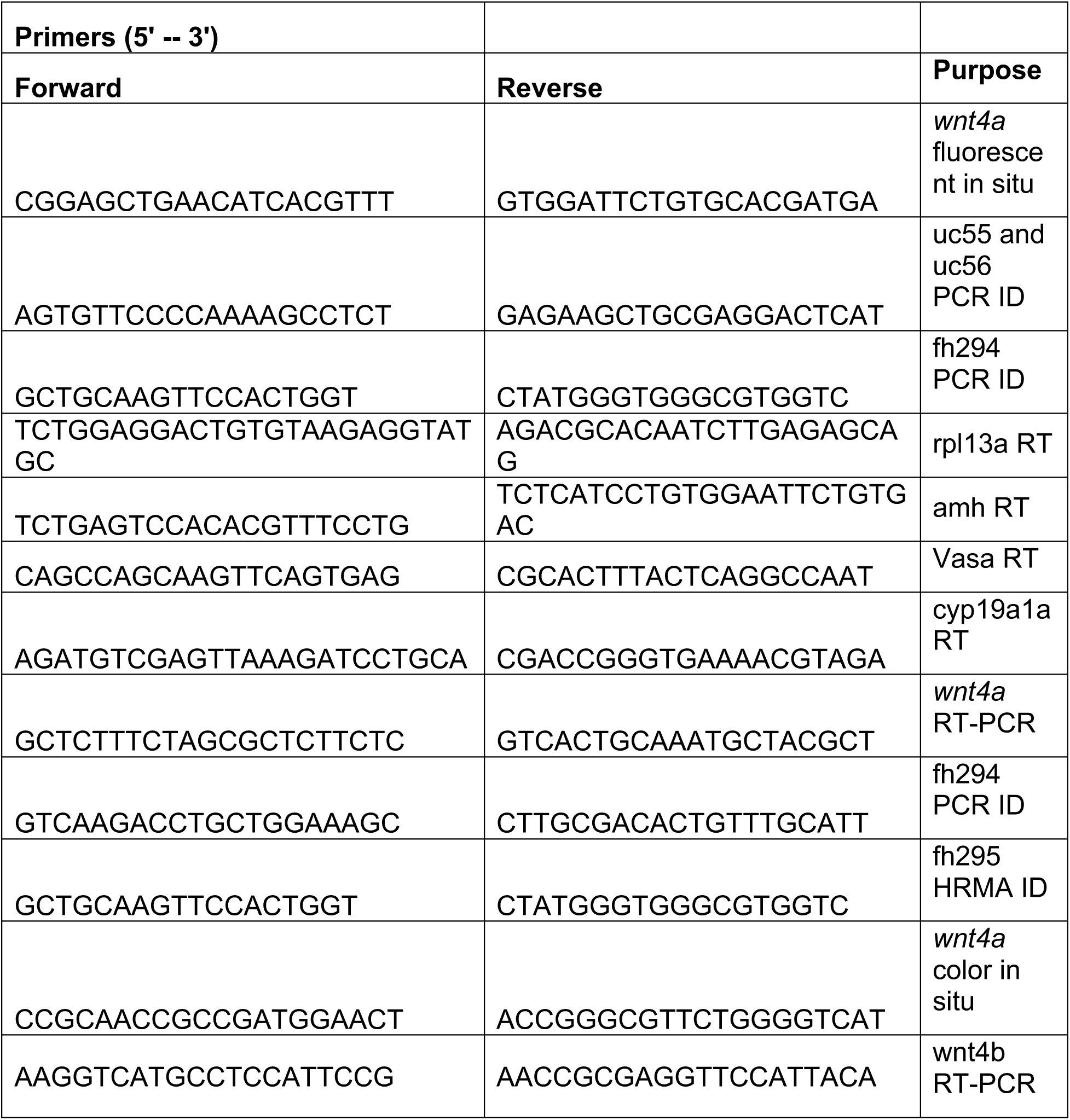
Oligos. Oligos listed were used in the excecution of this paper, 5’-→ 3’.

**Supplemental Table 2.** Genotypic ratios of *wnt4a* mutants were normal. The ratio of wild-type: heterozygous: mutant was not significantly different from expected Mendelian ratios, 1:2:1, consistent with normal survival.

|  | Number of Wild-types | Number of Heterozygotes | Number of Mutants | Resulting Ratio | Chi-square test |
| --- | --- | --- | --- | --- | --- |
| **fh295** | 56 | 124 | 51 | 1 : 2.21 : 0.91 | P = 0.48 |
| **fh295** | 162 | 307 | 143 | 1 : 1.90 : 0.88 | P = 0.55 |
| **uc55** | 103 | 202 | 119 | 1 : 1.96 : 1.16 | P = 0.34 |
| **uc56** | 109 | 208 | 79 | 1 : 1.91 : 0.72 | P = 0.06 |

## SUPPLEMETNAL FIGURE LEGENDS

**Supplemental Figure 1. In zebrafish gonads, *wnt4a* expression is female specific.** RT-PCR analysis of gene expression in adult (90dpf) (A) and juvenile (30dpf) (B) gonads. The germ cell-specific gene *vasa* was detected in both ovaries and testes. The female-specific gene *cyp19a1a* was detected in only ovaries while the male-enriched gene *amh* was detected at slightly higher levels in testes compared to ovaries. Expression of *wnt4a* was only detected in ovaries, while *wnt4b* transcript appeared faintly in the testies. The ribosomal protein gene *rpl13a* is ubiquitously expressed used here as loading control. NRT, no reverse transcriptase control.

**Supplemental Figure 2. *wnt4a* is expressed in somatic cells that surround stage 1b oocytes in 23dpf female zebrafish.** A. Confocal micrographs showing *wnt4a* RNA-expressing somatic gonadal cells (red) are closely associated with Vasa-expressing (green) oocytes. Wnt4a does not appear to be expressed in theca cells, which express Tg(*cyp19a1a:egfp*) (white, n=3). Red staining that overlaps with white was consistent with the background staining see throughout the tissue. B. In a simplified magnified view outlined in A, *wnt4a* localizes to the somatic cells adjacent to an oocyte with a diameter of 23µm. Expression of *wnt4a* is found consistently in somatic cells of stage 1b oocytes that are >20µm in diameter. Nuclei are stained blue. Scale bar in A, 10µm).

**Supplemental Figure 3. CRISPR/Cas9 mutant allele generation**. *wnt4a(uc55)* and *wnt4a(uc56)* were both generated by targeting the highlighted sequence (red) of Exon two. Mutagenesis resulted in 17bp (uc55) or 23bp (uc56) insertions (bold), PAM sequence (underlined).

**Supplemental Figure 4. *wnt4a* mutant fish have normal secondary sexual characteristics.** Light micrograph pictures showing that the pigment patterns and body shapes of *wnt4a(uc55)* mutant females and males (B-B’” and D-D’”, respectively) are indistinguishable from their wild-type female and male siblings (A-A’” and C-C’”, respectively). Panels A-D compare of body shapes. B’-D’ and B”-D” are low and high magnification views, respectively, of anal fin pigmentation. Note that males have more yellow pigmentation than females. A’”-D’” show high magnification views of the genital orifice. Note that the genital orifice of females protrudes from the ventral body wall, while those of male do not (black arrow).

**Supplemental Figure 5. *wnt4a* mutant zebrafish have normal mature gonads.** Wild-type males (A) and *wnt4a* mutant males (B) at 90dpf both have all stages of spermatogenesis and are indistinguishable. Similarly, wild-type females (C) and *wnt4a* mutant females (D) have all stages of oogenesis and are indistinguishable. Scale: 1mm

**Supplemental Figure 6. 3D renderings of the male reproductive ducts.** WT male (left) develop a fused duct deferens which is absent in the mutant (right).

**Supplemental Figure 7. 3D renderings of the female reproductive ducts.** WT female (left) develop a complete oviduct which fails to connect to the genital orifice in the mutant (right).

**Supplemental Figure 8. Growth of the male zebrafish reproductive duct over time.** Males were measured for duct elongation as a percentage of the distance between the posterior end of the testis and the genital orifice. Males were collected from 25dpf through 55dpf including adults and elderly (about 2 years old) for reference. Wild-type and *wnt4a* heterozygous males (green) had fully elongated reproductive ducts by 55dpf, while ducts in *wnt4a* mutants (yellow) never fully connected to the genital orifice.

